# Benchmarking organ-specific responses to therapies in tissues differentiated from Cystic Fibrosis patient derived iPSCs

**DOI:** 10.1101/2024.03.13.584768

**Authors:** Abdelkader Daoud, Sunny Xia, Onofrio Laselva, Janet Jiang, Christine E. Bear

## Abstract

Cystic Fibrosis (CF) is a life-shortening disease that is caused by mutations in the *CFTR* gene, a gene that is expressed in multiple organs. There are several primary tissue models of CF disease, including nasal epithelial cultures and rectal organoids, that are effective in reporting the potential efficacy of mutation-targeted therapies called CFTR modulators. However, there is the well-documented variation in tissue dependent, therapeutic response amongst CF patients, even those with the same CF-causing mutation. Hence, there is an interest in developing strategies for benchmarking therapeutic efficacy in different organs relative to isogenic controls. In this study, we evaluated the CFTR chloride channel response to the highly effective CFTR modulator: Trikafta, in CF patient specific, iPSC-derived colonic and airway cultures relative to mutation-corrected (non-CF) tissues from that same individual. We measured pharmacological rescue in both tissues, but interestingly, Trikafta treatment resulted in different levels of functional rescue in the two tissues relative to the relevant isogenic control. This proof-of-concept study lays the groundwork for future comparisons of patient-specific CF therapeutic responses in both pulmonary and extra-pulmonary systems.

## Introduction

### Cystic fibrosis

(CF) is an autosomal recessive disease caused by mutations in the *Cystic Fibrosis Transmembrane Conductance Regulator* (*CFTR)* gene. CF affects multiple organs including the lungs, pancreas, liver, intestine and the reproductive tract. To date, there are more than 700-disease causing mutations in the CFTR gene. Encouragingly, people harbouring certain mutations, including the most common mutation F508del, have access to a highly-effective therapy called Trikafta. This drug contains three small molecules, i.e., two corrector compounds and one potentiator compound. The two correctors, tezacaftor and elexacaftor act to rescue the basic defects in F508del-CFTR protein folding, assembly and trafficking through the cellular biosynthetic compartment. The potentiator compound, ivacaftor, augments the open probability of the cyclic AMP-activated F508del-CFTR channel. Clinical use of Trikafta (ETI) can result in substantial improvements in lung function with reductions in respiratory symptoms and bacterial exacerbations in most people with CF harbouring at least a single copy of F508del (1, 2). However, the impact of Trikafta on CF-related disease in other organs, including the colon remains unclear (3, 4). Unfortunately, extra-pulmonary disease in CF can exert a major toll on the quality of life of a significant number of people with this disease. Hence, there is an unmet need for CF-patient specific tissue models that report the consequences of disease-causing mutations and therapeutic interventions in both pulmonary and extrapulmonary systems.

In this study, we leveraged a fluorescence-based assay of the functional rescue of mutant CFTR (i.e, F508del-CFTR) (5) to develop a strategy for benchmarking tissue dependent, patient specific drug responses. This method enables direct comparison of the functional rescue of F508del-CFTR channel activity in the apical membrane of 2D colonic and lung tissues differentiated from patient specific iPSCs in a medium-throughout manner. Comparative studies with lung and colonic tissues differentiated from mutation corrected iPSCs derived from the same individual permitted evaluation of the relative, tissue dependent, functional rescue by Trikafta.

## Materials and methods

### 2.1 Generation and maintenance of human induced pluripotent stem cells (iPSCs)

Human induced pluripotent stem cells (iPSCs) were generated as previously described (6) by the Centre for Commercialization of Regenerative Medicine (CCRM) for the CFIT program donors. Peripheral blood mononuclear cells (PBMCs) from donor # 1 and donor #2 were isolated and reprogrammed into iPSCs using the Sendai Reprogramming Kit (Thermo Fisher, cat # A34546). CFLine#1 was generated from donor #1 while CFLine#2a and CFLine#2b were derived from donor # 2. The isogeneic control line (WtLine#1) was generated from CFLine#1 using the GeneArt™ CRISPR Nuclease Vector with CD4 Enrichment Kit (Thermo Fisher, cat # A21175); The genetic correction of F508del in CFTR gene was verified by CCRM using PCR and Sanger sequencing at the targeted locus. iPSC lines pluripotency was determined by CCRM using flow cytometry and qRT-PCR; flow cytometry showed that >80% of the population was positive for surface markers SSEA4, TRA-1-60 and intracellular markers OCT4 and SOX2. qRT-PCR revealed that pluripotency associated genes including OCT4, NANOG and DNMT3B were expressed at >80% in comparison to an hESC reference line. iPSCs were plated on Matrigel® hESC-Qualified Matrix (Corning Inc., cat # 354277) coated 6-well plate in mTeSR1 (STEMCELL Technologies, cat # 85870) media and were passaged every 4 days at a ratio of 1:6 t o1:8. For cryopreservation, cells grown in a 6-well plate were collected, centrifuged at 300g for 3min and resuspended in mFreSR (STEMCELL Technologies, cat # 05855); iPSCs from each well of the 6-well plate were stored per vial and placed in liquid nitrogen until ready to thaw. Cells were thawed into Matrigel® hESC-Qualified Matrix-coated 6-well plates. In general, one vial of cells cryopreserved as described above was thawed into 1-2 wells of the 6-well plate. CFline#1 were received from CCRM at passage P3+6 and WtLine#1 was at P3+33. CFline#2a and CFLine#2b were both at passage P3+9. Passages used in the study were between 1 and 6 passages from the stock vials. All lines tested negative for mycoplasma using Lonza MycoAlert Plus kit (Lonza, cat # LT07-710).

### 2.2 Generation and maintenance of Human colonic organoids (HCOs)

iPSCs were plated on Matrigel® hESC-Qualified Matrix (Corning Inc., cat # 354277) coated 24-well plate in mTeSR1 (STEMCELL Technologies, cat # 85870) media supplemented with 10μM of the ROCK inhibitor Y27632 (STEMCELL Technologies, cat # 72302). 24h later, medium was changed to mTeSR without Y27632 and cells were incubated for an additional 24h in a 37 °C, 5% CO2 incubator. The next day, iPSCs were incubated with the definitive endoderm (DE) induction media containing 100 ng/mL of Activin A (Cell guidance Systems, cat # GFH6-100x10) for 3 days. Next, the DE monolayer was incubated for 4 days with the hindgut induction medium supplemented with 500 ng/mL of FGF4 (R&D Systems, cat # 235-F4-025/CF) and 3μM of CHIR99021 (STEMCELL Technologies, cat # 72052) to induce the formation of mid/hindgut endoderm (MHE). Uniform clumps collected from the MHE monolayer are mixed with Matrigel® Basement Membrane Matrix (Corning Inc. cat # 354234) and plated in a 24-well plate as 50uL Matrigel® bubbles. The immature structures are patterned into colonic organoids by incubation in an intestinal growth medium (Advanced DMEM/F-12, N2, B27, 15 mM HEPES, 2 mM L-glutamine, penicillin-streptomycin) supplemented with 100 ng/mL EGF (R&D Systems, cat # 236-EG) and 100 ng/mL of BMP2 (R&D Systems, cat # 355-BM-100) for 3 days. Media are then changed to the intestinal growth medium supplemented with 100 ng/mL EGF alone for 11 days. On day 21, human colonic organoids at passage 1 (OP-1) are split at 1:2 into a new 24-well plate (OP-2) and are incubated in a previously described growth factor conditioned media (7, 8). Media are changed every 2-3 days until day 35, when HCOs are split every 7-8 days for functional studies. HCOs were tested for CFTR function at passages between 1 and 6. In all experiments, at least 3 biological replicates (organoids passages) were tested in each study with at least 4 technical replicates.

### 2.3 Immunofluorescence Staining

Organoids were fixed in 4% PFA and embedded in Tissue-Tek® O.C.T (VWR, cat # 25608-930). Slides of cryosectioned organoids were blocked in 5% normal donkey serum (Jackson Immuno Research, cat # AB_2337258) for 30 min at room temperature. Next, the samples were incubated with primary antibodies overnight at 4°C. Following three washes in PBS-0.5% Triton-X (PBST), samples were incubated with the secondary antibodies and DAPI for 2h at room temperature. Slides were then washed twice with PBS-T, once with PBS and mounted using Fluoromount-G® (SouthernBiotech, cat # 0100-01). Confocal images were taken using a Zeiss LSM 880 NLO with Airyscan confocal microscope.

### 2.4 The Apical Chloride Conductance (ACC) Assay

The Apical Chloride Conductance (ACC) Assay for the measurement of CFTR Function was performed following the protocol described by Xia *et al*. (8) and Ahmadi *et al*. (9). Briefly, HCOs were removed from the Matrigel® Basement Membrane Matrix (Corning Inc. cat # 354234) bubble, collected in ice-cold DMEM/F12 and pelleted through centrifugation at 500× g for 5 min. The organoids were then re-suspended in growth factor conditioned media and plated onto 0.01% Poly-L-Lysine (Sigma-Aldrich, cat # P4707) coated 96-well plates (Corning cat # 3603). 24h later, the medium was changed with fresh growth factor conditioned media and opened organoids were treated with different CFTR modulators. Organoids reached a confluence of 80% after 24h post plating and almost 100% on the day CFTR conductance assay was performed. The ACC assay was always performed two days post organoids plating. In this assay, HCOs were incubated with zero sodium, chloride and bicarbonate buffer containing 0.5 mg/mL of FLIPR® dye (Molecular Devices, cat # R8042) for 30 min at 37ºC. The 96-well plate containing the organoids was then loaded into the FLIPR® Tetra station (Molecular Devices) and simultaneous images of the entire plate were acquired. Data analysis was performed following steps described previously (8).

### 2.5 Western blotting

After the ACC assay was performed, the FLIPR dye was removed from the wells, and the plates were stored at -20°C. In general, multiple plates were lysed simultaneously to avoid batch to batch differences. To lyse the organoids, 50uL of lysis buffer (50 mM Tris-HCl, 150 mM NaCl, 1 mM EDTA, pH 7.4, 0.2% (v/v) SDS, 0.1% (v/v) Triton X-100, 1X inhibitor cocktail) was added per well and the plates were placed on ice with gently rocking for 10min. After centrifugation at 13,00orpm for 10 min at 4 °C, 40uL of lysate was mixed with 5X Laemmli Sample Buffer and loaded in each well in a 6% Tris-Glycine gel. The gel was then transferred to a nitrocellulose membrane on ice for 1 hour at 100V. The membrane was blocked in 5% milk for 30min and incubated with mouse monoclonal CFTR 596 (1:1000, UNC CFTR antibodies) antibody, and rabbit Calnexin (1:5000, Sigma-Aldrich, cat # C4731) antibody overnight at 4°C. Following three washes with PBS-0.1%Tween 20, the blots were further incubated with the corresponding secondary anti-mouse HRP (1:5000, Abcam, ab6789) and anti-rabbit HRP (1:5000, Abcam, ab6721) antibodies for 1 hour at room temperature. Finally, the immunoblots were washed three times with PBS-0.1%Tween 20 and developed on the Li-Cor Odyssey Fc (LI-COR Biosciences) station.

### 2.6 Statistical Analysis

Unpaired two-tailed t test was performed on data with two datasets. Ordinary one-way ANOVA with Tukey’s multiple comparison test was performed on all data with more than two datasets. Correlations between datasets were assessed using the “Spearman’s test”. p < 0.05 was considered statistically significant. Statistical analyses were performed using GraphPad Prism 10.10.

## Results

### 3.1 2D Colonic organoids exhibit CFTR mediated apical chloride conductance

3D colonic organoids were generated from an iPSC line harbouring Wt-CFTR using a previously established protocol (10, 11) (Figure 1A). Immunostaining confirmed the expression of CDX2, an intestinal marker, and SATB2, a colonic marker, within the HCOs (Figure 1B). To measure the apical conductance mediated by CFTR, 3D HCOs were converted into a 2D-format to facilitate the splitting open of organoids, exposing the apical membrane (Figure 1C). The method for “opening the organoids” was previously published (8, 12) and is described in the methods section (2.3 The Apical Chloride Conductance (ACC) Assay).

**Figure 1:**
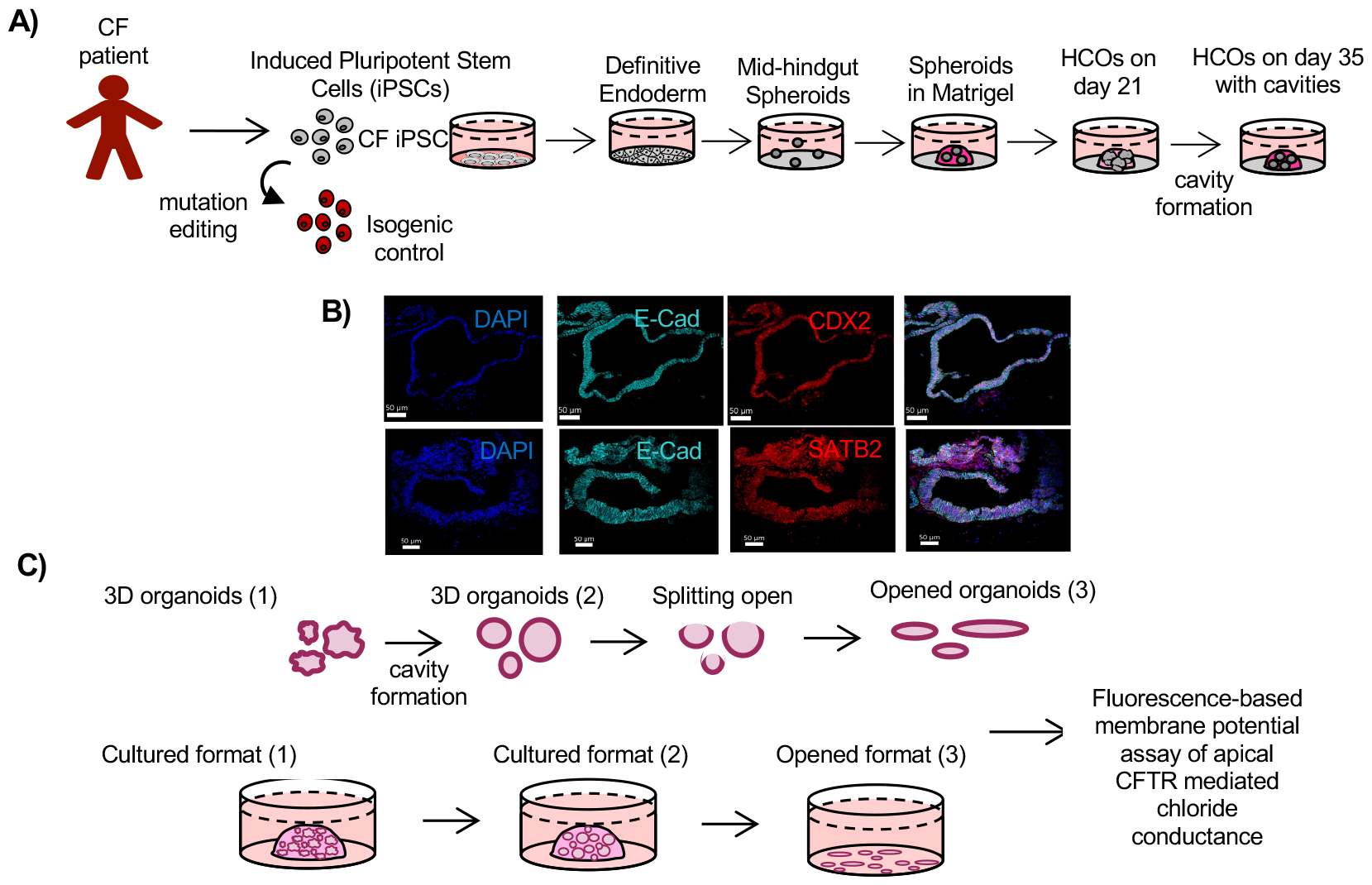
**A)** General protocol for generating human colonic organoids (HCOs) as previously described (10, 11), from patient-derived iPSCs. **B)** HCO stained for epithelial marker E-cadherin, intestinal epithelial marker CDX2, colonic marker SATB2 and DAPI (bottom). Images were obtained 20x magnification. Scale bar = 50 μm. **C)** iPSC-derived spheroids split open as described by Xia *et al*. (8) and Ahmadi *et al*. (9) for subsequent fluorescence-based assay of membrane potential (FMP) and apical chloride conductance.

The fluorescence membrane potential (FMP) assay, that we modified to measure CFTR function in epithelial tissues (also known as the apical CFTR-mediated chloride conductance assay: ACC), (5, 8, 9, 13), was used to confirm the functional expression of CFTR in 2D colonic organoids (Figure 2A). Sensitivity of the fluorescence response to the inhibitor: CFTRInh-172 supported our claim that the fluorescence that it was mediated by CFTR (Figure 2A). To validate the reproducibility of the assay, we compared vehicle (DMSO) versus forskolin stimulated CFTR function across multiple organoid passages, ranging from 3 to 6 in the non-CF iPSC line (WtLine#1), (Figure 2B).

**Figure 2:**
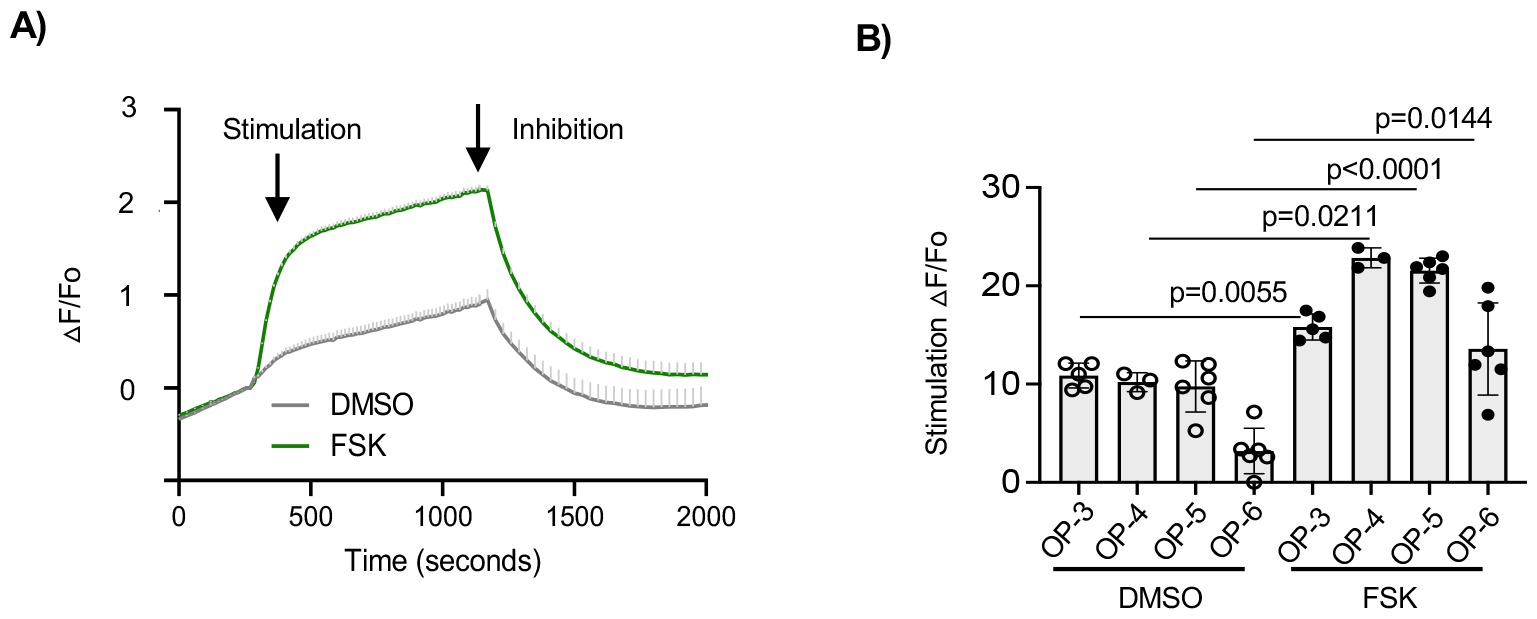
FMP assay of Wt-CFTR channel activity in split open colonic organoids generated from non-CF iPSC line (WtLine#1). **A)** sample FMP trace showing activation and inhibition of CFTR channel activity in HCOs from organoid passage 5 (OP-5). **B)** Bar graph showing passage dependent, CFTR-mediated change in FMP response after stimulation by FSK (statistical significance was tested using one-way ANOVA by comparing FSK groups to DMSO conditions).

### 3.2 2D CF Colonic organoids exhibit functional rescue of mutant F508del-CFTR after treatment with the CFTR Modulator TRIKAFTA

Multiple passages of HCOs (passages 3-6) were generated from CFline#2a, an iPSC line derived from a CF patient carrying the F508del mutation (Figure 3A). In order to determine the extent of functional rescue with TRIKAFTA (ETI), which includes VX-661 (3µM), VX-445 (3µM), and VX-770 (1µM) (1, 2), the organoids were opened to generate the 2D format as previously described. The 2D organoids were treated for 24 hours with the corrector compounds: VX-661 and VX-445 or the vehicle (DMSO) as the control. Next, the 2D organoids were plated for study using the FMP assay and responses to acute treatment with forskolin and the potentiator (VX-770), measured. As evident in figure 3A and 3B, pre-treatment with the corrector compounds augmented the response to forskolin and VX-770 as expected. There was a significant improvement in functional expression of F508del-CFTR after correction with VX-661 and VX-445 relative to DMSO-pretreated 2D cultures in the 4 different passages tested. Western blot analyses of F508del-CFTR expression, confirmed that the corrector compounds were effective in rescuing the misprocessing defect induced by the F508del mutation in HCOs generated from this patient derived iPSC line (Figure 3C). Specifically, mature CFTR protein (band C) in HCOs passages 3-6, appeared following 24-hour treatment with VX-661 plus VX-445 as expected. As shown in the supplementary data, the rescue effect of the Trikafta combination was replicated using a second iPSC line from the same individual (CFline#2b) donor and in a line from a unique donor (CFline#1) in passages 3 and 6.

**Figure 3:**
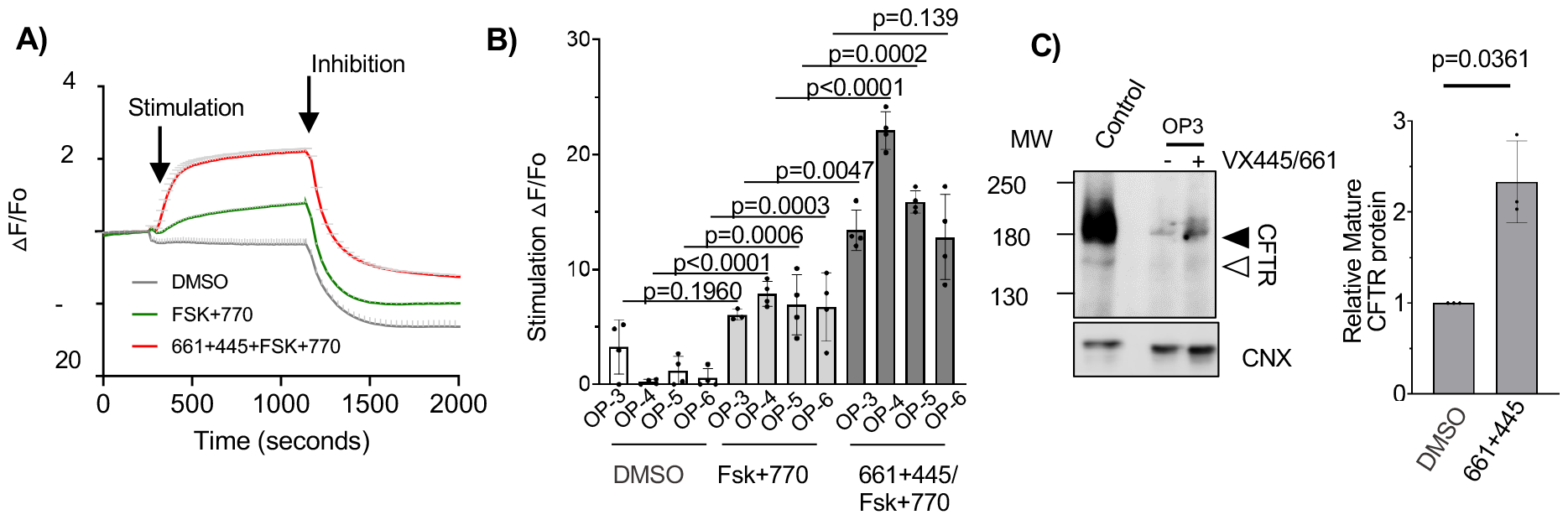
FMP assay of mutant CFTR channel activity in CF (genotype= F508del/F508del) colonic organoids (OP-4) provides reproducible measurements of drug response. **A)** A sample FMP trace showing activation and inhibition where cultures were pre-treated and acutely treated with vehicle alone (DMSO: grey line), or pretreated with DMSO and acutely activated with forskolin (FSK) and the potentiator VX-770, (FSK+770: green line) or pretreated with correctors VX-661 plus VX-445 and acutely activated with forskolin and VX-770, (661+445+FSK+770: red line). **B)** Bar graph summarizes responses for studies in A with reproducibility of technical replicates shown (3 wells of 96 well plate). Significant responses were measured in iPSC-derived HCOs from dF508 CF patient (CFline#2a) at four different organoid passages except in conditions indicated for passage 3 and passage **6**. Statistical significance tested using one-way ANOVA. **C)** Western blotting of CFline#2a organoids for CFTR and Calnexin showing an increase in CFTR band C after drug rescue with VX-445+VX-661 in organoid passage 3. The control is a lysate from non-CF primary rectal organoids. The adjacent bar graphs shows that the abundance of band C (the mature form of CFTR protein) is augmented by the correctors VX-445+VX-661 as expected in 3 different organoid passages.

### 3.3 Functional Rescue of 2D CF Colonic organoids and 2D CF immature lung tissue by Trikafta, relative to patient-specific iPSC line corrected to Wt-CFTR

The degree of functional rescue of F508del-CFTR in colonic or lung tissues was determined relative to CFTR mutation corrected tissues from the same individual. In figure 4A, we show that Trikafta (ETI) treated 2D colonic organoids exhibited improvements in F508del-CFTR mediated channel function up to values that were comparable to function measured in mutation-corrected colonic tissues from the same individuals. Interestingly, the functional rescue mediated by Trikafta in immature lung cultures, (generated as previously described,(5)), only partially recued F508del-CFTR function (Figure 4B). There remained a significant difference between the CFTR function measured in mutation corrected lung tissues and pharmacologically corrected tissue.

**Figure 4:**
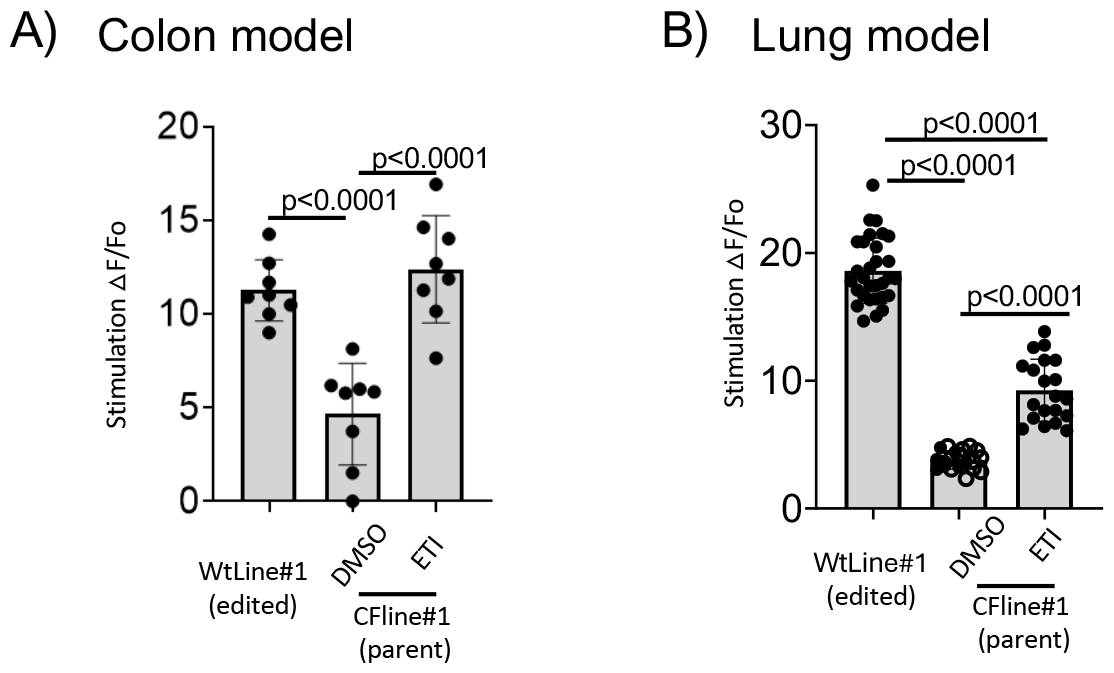
The effect of drug rescue versus genetic correction on CFTR function in colonic and lung tissues using the FMP assay. **A)** Bar graph showing CFTR partial rescue in colonic organoids from dF508 CF patient CFline#1 by TRIKAFTA and the isogenic control WtLine#1 by genetic correction in an FMP assay (data generated from two biological replicates each with 4 technical replicates; Statistical significance was tested using one-way ANOVA). **B)** Bar graph showing CFTR partial rescue in immature lung tissue from dF508 CF patient CFline#1 by TRIKAFTA and the isogenic control WtLine#1 by genetic correction in an FMP assay (Statistical significance was tested using one-way ANOVA).

## 4. Discussion

We described a strategy where the therapeutic efficacy of a CFTR targeted intervention can be tested in two different, organ-specific tissues differentiated from CF iPSCs and the response benchmarked relative to their non-disease, isogenic control tissue. The results of this comparative study suggest that the efficacy of Trikafta varies between these two tissue types, the colon and the airways. This approach has the potential to be applied toward evaluating therapeutic efficacy in many of the organ systems affected by Cystic Fibrosis, i.e, pancreatic and bile ductules and epididymis (14-17)

Given the current uncertainty regarding the response in different CF-affected organs to CFTR modulators and its potential toxicity, there is an urgent need to develop robust CF organ-specific platforms for drug testing. To date, there has been impressive progress in testing CFTR modulators in primary tissue cultures modeling the nasal, airway and rectal epithelium. Gonska and colleagues, showed that Trikafta was effective in rescuing the CFTR mediated chloride conductance across the apical membrane of patient derived nasal and rectal epithelial cultures (18). In these studies of tissue from patients harbouring F508del and rare mutations, the rescue of CFTR conductance was measured in rectal tissue in both 2D and 3D models using bioelectric methods or imaging of CFTR-mediated spheroid swelling respectively. Altogether, the positive responses in both tissues, supported the claim that Trikafta was effective in the airways and in the colon of people bearing the major mutation or certain rare mutations. The group of Cutting and colleagues, also employed a multimodal approach to examine the efficacy of Trikafta in an ultra-rare CF-causing mutation (19). The positive response measured in 2D and 3D cultures, modeling airway and rectal epithelia, supported the potential efficacy in Trikafta in treating disease in both organs in people with this ultra-rare mutation. Although comprehensive, both studies lacked a strategy for benchmarking organ-specific response within each individual. Without a patient-specific control, it is problematic to attribute relatively poor responses to a lack of therapeutic efficacy rather than poor model quality.

The 2D tissue models employed in this study are considered relatively immature. We characterized the airway tissue generated while submerged in a 2D format in our previously published work (5). While the functional expression of CFTR is relatively high in these cultures, transcriptomic analyses support the claim that the tissue models the lung progenitor stage. It has been shown that iPSC-derived HCOs contain colonic MUC5b-expressing goblet cells and exhibit increased numbers of MUC2 goblet cells by day 35 (Munera, 2017 # 1, Daoud, 2020 # 2). Despite the fact that at this stage, several other tissue-specific cell types are absent, HCOs demonstrate competence in producing other cells such as enteroendocrine cells after *in vivo* transplantation into immunodeficient mice.

In our knowledge, this is the first report of the use of isogenic controls to benchmark organ-specific responses to CFTR targeted therapies. However, there are experimental caveats associated with this study. The differentiation protocols employed lead to the generation of relatively immature colonic or airway tissues. These protocols were chosen because they lead to the production of tissue expressing CFTR protein in a 2D format suitable for study of apical, CFTR mediated chloride conductance using the medium throughput fluorescence-based assay of membrane potential changes. This assay enables simultaneous measurements of replicates samples optimizing confidence in the results. However, the relatively immature tissues do not contain all of the cell types of the mature epithelium (5). For example, the immature lung cultures do not express the ionocyte, a rare cell type proposed to have a major role in fluid transport. It remains possible that the differential responses to Trikafta observed in the iPSC - derived colonic and airway tissues are related to their differentiation status. Future studies are required to evaluate Trikafta responses in airway and colonic tissues that are sufficiently differentiated to express all of the cell types know to populate the mature tissue.

## Acknowledgements

The authors acknowledge the assistance by the program managers of the CFIT program (Tarini Gunawardena and Paul D.W. Eckford) in identifying demographic data for the CFIT tissue donors. We also acknowledge funding by the Cystic Fibrosis Canada and the Hospital for Sick Children.

## Figure Legends

**Supplementary Figure:**
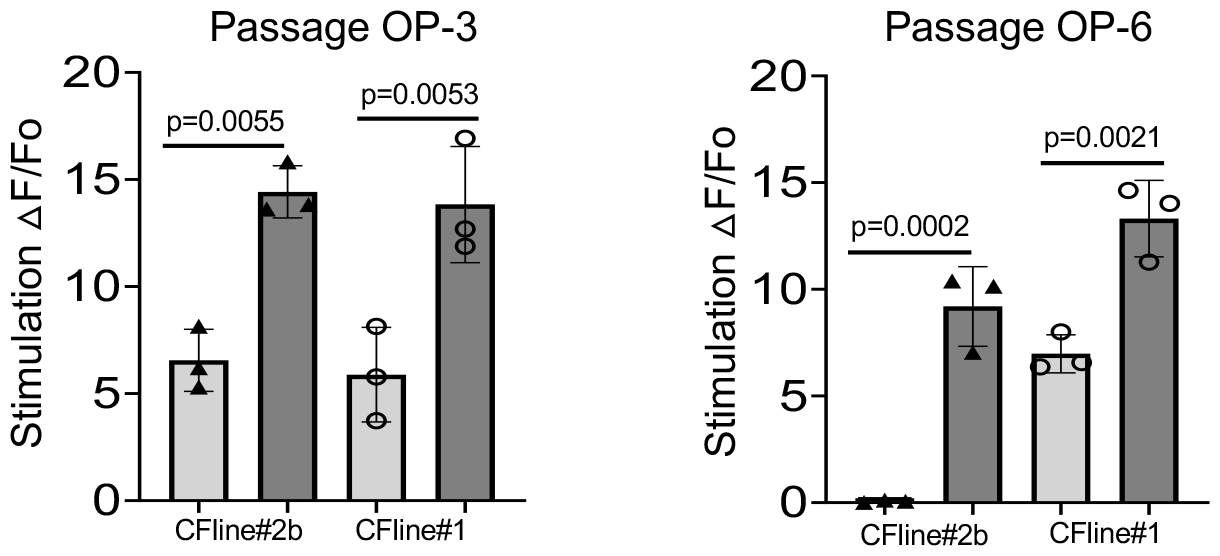
FMP assay of CFTR channel activity in CF colonic organoids provides reproducible measurements of drug response. Bar graphs showing rescue of CFTR channel activation in F508del/F508del colonic organoids from two passages (OP-3 and OP-6) of two lines CFline#2b and CFline#1 after TRIKAFTA correction in an FMP assay (Statistical significance was tested using one-way ANOVA).

## Notes

### Competing Interest Statement

The authors have declared no competing interest.

